# GToTree: a user-friendly workflow for phylogenomics

**DOI:** 10.1101/512491

**Authors:** Michael D. Lee

## Abstract

**Summary:** Genome-level evolutionary inference (i.e., phylogenomics) is becoming an increasingly essential step in many biologists’ work - such as in the characterization of newly recovered genomes, or in leveraging available reference genomes to guide evolutionary questions. Accordingly, there are several tools available for the major steps in a phylogenomics workflow. But for the biologist whose main focus is not bioinformatics, much of the computational work required - such as accessing genomic data on large scales, integrating genomes from different file formats, performing required filtering, stitching different tools together, etc. - can be prohibitive. Here I introduce GToTree, a command-line tool that can take any combination of fasta files, GenBank files, and/or NCBI assembly accessions as input and outputs an alignment file, estimates of genome completeness and redundancy, and a phylogenomic tree based on the specified singlecopy gene (SCG) set. While GToTree can work with any custom hidden Markov Models (HMMs), also included are 13 newly generated SCG-set HMMs for different lineages and levels of resolution, built based on searches of ~12,000 bacterial and archaeal high-quality genomes. GToTree aims to give more researchers the capability to make phylogenomic trees.

**Availability:** GToTree is open-source and freely available for download from: github.com/AstrobioMike/GToTree

**Documentation:** github.com/AstrobioMike/GToTree/wiki

**Implementation:** GToTree is implemented primarily in bash, with helper scripts written in Python.

## 1 Introduction

The number of sequenced genomes is increasing rapidly, largely through the recovery of metagenome-assembled genomes (MAGs) (e.g. Hug et al. 2016; Parks et al. 2017) and through the generation of single-cell amplified genomes (SAGs) (e.g. Kashtan et al. 2014; Berube et al. 2018). Phylogenomics (inferring genome-level evolutionary relationships) is becoming a fundamental step in the characterization of newly recovered genomes. Additionally, large-scale comparative genomics efforts leveraging growing public databases can be employed to investigate evolutionary avenues such as ancestral reconstruction (Braakman, Follows, and Chisholm 2017), which are guided by phylogenomics.

There are several tools available for the major steps in a phylogenomics workflow, and at least one analysis platform that incorporates a phylogenomics workflow amid a larger infrastructure (anvi'o; Eren et al. 2015). But a complete workflow focused solely on phylogenomics, enabling greater efficiency and scalability, and with flexibility with regard to input formats, is lacking.

GToTree fills a void on three primary fronts: 1) it accepts as input any combination of fasta files, GenBank files, and/or NCBI accessions - allowing integration of genomes from various sources and stages of analysis without any computational burden to the user; 2) it enables the automation of required between-tool tasks such as filtering out hits by gene-length, filtering out genomes with too few hits to the specified target genes, and swapping genome identifiers so resulting trees and alignments can be explored more easily; and 3) its scalability - GToTree can turn ~1,700 input genomes into a tree in one hour on a standard laptop, and can optionally run many steps in parallel. This software gives more researchers the capability to create phylogenomic trees to aid in their work.

## 2 Description

### 2.1 Input

The required inputs to GToTree are 1) any combination of fasta files, GenBank files, and/or NCBI assembly accessions, and 2) an HMM file with the target genes. The HMM file can be custom or one of the 13 included HMM files covering varying breadths of diversity. Optionally, the user can also provide a mapping file of specific input genome IDs with the labels they would like to have displayed in the final alignment and tree.

### 2.2 Processing

An overview of the GToTree workflow is presented in Figure 1 and detailed here:

1. Retrieve coding-sequences (CDSs) for input genomes, depending on the input source:

- fasta files - identify CDSs with prodigal (Hyatt et al. 2010)
- GenBank files - extract CDSs if annotated, if not identify with prodigal (Hyatt et al. 2010)
- NCBI accession - download GenBank and RefSeq assembly summary files as needed, build appropriate ftp paths, attempt to download just the amino acid sequences of CDSs if annotated, if not, download the assembly in fasta format and identify CDSs with prodigal (Hyatt et al. 2010)
2. Identify target genes in all genomes with HMMER3 (Eddy 2011) using pre-defined model cutoffs (--cut_ga)
3. Report estimates of genome completeness/redundancy using the information from the HMM search
4. Filter gene-hits and genomes, add in gap-sequences:

- filter out gene-hits based on length (get the median of all genes in that set, filter out those whose length is not within a specified range of the median length)
- filter out genomes if they do not have successful hits to at least a specified fraction of the total target genes searched
- add in gap-sequences for target genes that are missing from genomes retained in the analysis
5. Align, trim, and concatenate:

- align each gene-set with Muscle (Edgar 2004)
- perform automated trimming with Trimal (Gutierrez et al.2009)
- concatenate all gene-sets together into full alignment
6. Optionally add more informative genome labels to be displayed in the alignment and tree, with either or both of the following:

- a two-or three-column tab-delimited mapping file with input genome ID in column 1, either the desired label in column 2 to swap entirely, and/or a string of text to append to the label in column 3 (not all input genomes need to be specified)
- use TaxonKit (Shen and Xiong 2019) in order to add lineage information to genome labels that had associated taxids (whether from NCBI accessions or found in the provided GenBank files)
7. Tree with FastTree (Price, Dehal, and Arkin 2010).

**Figure 1:**
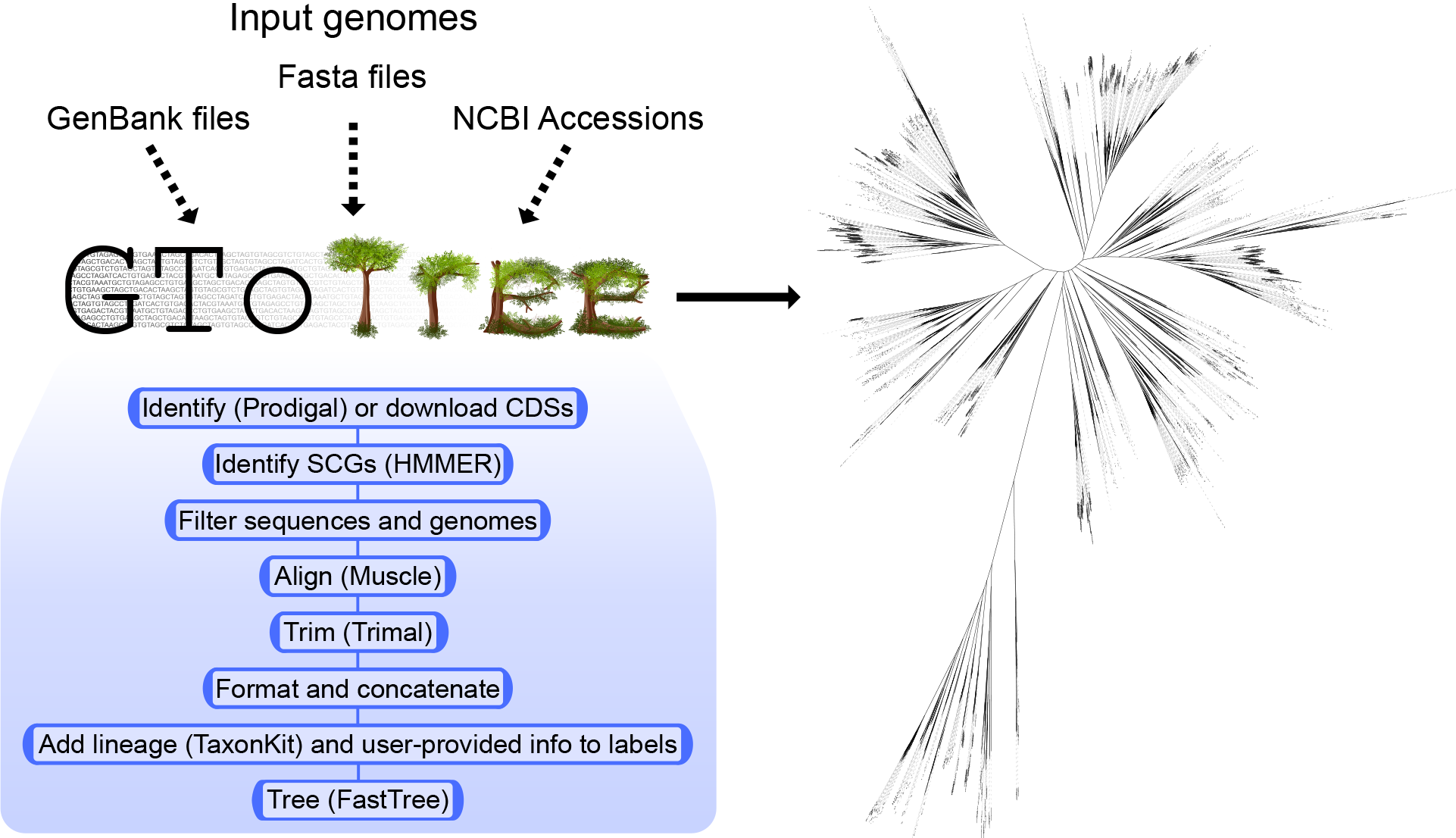
Overview of general workflow and an example Tree of Life made with GToTree encompassing ~1,700 genomes from NCBI's RefSeq using a universal SCG-set (Hug et al. 2016).

### 2.3 Outputs

The primary outputs from GToTree include the full alignment file (fasta), the tree file (newick), and tab-delimited summary tables with information on all genomes and individual ones for each genome input source. Additionally, outputs include report files on filtered or problematic genes/genomes.

### 2.4 SCG-set generation

All 17,929 Pfam (protein families; El-gebali et al. 2019) HMM profiles from release 32.0 (accessed on Dec. 2018) were downloaded from the Pfam ftp site (ftp://ftp.ebi.ac.uk/pub/databases/Pfam/). As Pfam-HMMs actually target specific domains or protein regions, there are many unique Pfam entries that come from the same functional protein - e.g. Enolase_N (PF03952) and Enolase_C (PF00113). This is not ideal if using them to search for SCGs for purposes such as phylogenomics or completion/redundancy estimates. To ensure no two Pfam-HMMs from the same protein were contained in a SCG-set, only Pfams with HMMs that on average covered greater than 50% of the underlying protein sequences that went into building that Pfam’s HMM were retained. This left 8,924 Pfams.

Amino-acid coding-sequences of all “complete” genomes with annotations in NCBI were downloaded for bacteria (N=11,405; accessed 09-Dec-2018) and archaea (N=309; accessed 15-Dec-2018). All protein sequences were searched against the 8,924 filtered Pfam-HMMs with ‘hmmsearch’ (HMMER v3.2.1; Eddy 2011) with default settings other than specifying the “--cut_ga” flag to utilize the gathering thresholds stored in the curated Pfam models. Reported protein hits for each individual Pfam were tallied for each individual genome (Supp. Table 1). SCG-sets were generated for all Bacteria, all Archaea, and then for each bacterial phylum that held greater than 99 genomes, and each proteobacterial class that had greater than 99 genomes. For each of those taxonomic groups, Pfams that had 1 hit in greater than or equal to 90% of the genomes of that group were retained as the SCG-set for that group. These are summarized in Supp. Table 2, and the code used to generate the bacterial set is presented at github.com/AstrobioMike/GToTree/wiki/SCG-sets.

## 3 Results

To exemplify GToTree, NCBI assembly accessions were downloaded for all RefSeq, complete, representative genomes (with the search query ‘ “latest refseq”[filter] AND “complete genome”[filter] AND “representative genome”[filter] AND all[filter] NOT anomalous[filter] ‘ performed on 20-Dec-2018). This resulted in 1,698 genomes spanning Archaea, Bacteria, and Eukarya. Using a SCG-set that spans all 3 domains (Hug et al. 2016), runtime to create this tree (Figure 1) was ~60 minutes on a standard laptop (used was a late 2013 MacBook Pro). The tree was visualized by uploading the output newick file to the web-hosted Interactive Tree of Life (Letunic and Bork 2016), all code to generate it and the results files come packaged with GToTree.

## Funding sources

This work was funded in part by NASA Space Biology under grant NNH16ZTT001 N-MOBE.

## Acknowledgements

I would like to sincerely thank Titus Brown, Craig Everroad, Arkadiy Garber, Elaina Graham, Joshua Kling, Gustavo Ramirez, and Nathan Walworth for their time and help with ideas, testing, and troubleshooting during development.

## Conflicts of interest

None declared.

